# Phenotypic plasticity and the anthropause: an urban bird becomes less aggressive

**DOI:** 10.1101/2022.09.12.507677

**Authors:** Marlene Walters, Eleanor S. Diamant, Felisha Wong, Christina Cen, Pamela J. Yeh

**Affiliations:** Department of Ecology and Evolutionary Biology, University of California, Los Angeles, CA 90095 USA; Santa Fe Institute, Santa Fe, NM 87501 USA

**Keywords:** anthropause, human disturbance, human activity, urbanization, territorial aggression, behavioral plasticity, phenotypic plasticity, stressor

## Abstract

Urban areas often impose strong, novel selection pressures on wildlife. Phenotypic plasticity is an important mechanism helping organisms establish populations in novel environments. Phenotypic plasticity can be difficult to study in urban wildlife because many urban environmental variables are challenging to isolate and manipulate experimentally. We took advantage of the COVID-19 lockdowns to assess whether urban birds expressed aggression differently when relieved from frequent encounters with humans. We measured the territorial aggression responses of resident dark-eyed juncos *(Junco hyemalis)* on an urban college campus in Los Angeles, USA. We found that the population overall displayed significantly reduced aggression in pandemic year 2021 compared to the typical year 2019. Furthermore, individuals measured in both 2019 and 2021 showed significantly reduced aggression during 2021, demonstrating that individual birds maintain phenotypic plasticity in this trait. Our results show that human disturbance likely has a significant effect on the aggressive behavior of urban birds.

## Introduction

Cities are novel habitats that present new stressors for wildlife to overcome. Urban landscapes often have high concentrations of paved surfaces and artificial structures (1), light, noise, and chemical pollution (2), and abundant exotic wildlife and vegetation, including unfamiliar predators (3). Crucially, urban wildlife must be able to tolerate high frequencies of human disturbance (4–8). Any of these factors can disrupt a species’ ability to acquire resources, communicate, or reproduce, thus limiting which species can successfully establish populations in cities (9–14). For these reasons, rapid urban expansion into natural areas is one of today’s leading threats to biodiversity (3,15,16,12–14,17).

Yet, some native species still persist in cities around the world (11,12,14). These urban populations frequently express modified behaviors compared to their wildland counterparts (18,11,19). Songbirds, which are well-studied in the context of urban adaptation, show a range of modifications, including reduced fear of humans (20–22), use of artificial structures and materials in nesting (23,24), extended breeding seasons that in turn produce more offspring (25,26), modified long-range vocalizations (27), and modified territorial responses (28,29). These behavioral shifts typically represent examples of phenotypic plasticity, which is widely considered a key mechanism allowing species to survive in novel environments (30–34,11).

Measuring behavioral plasticity in urban wildlife is challenging because urban environmental variables are difficult to isolate and test experimentally on free-living populations. In particular, the amount of human activity in a given location is one urban variable that is especially difficult to control. Thus, the period of city-wide lockdowns in response to the COVID-19 pandemic, since termed the “anthropause” (35), provided a unique opportunity to observe free-living individuals in their home environments while levels of human disturbance were drastically altered (35–37). Recent studies from the anthropause show that urban bird populations demonstrated significant differences in communication (36,38) and nesting behavior (39) while environmental stressors were temporarily relaxed.

We used the natural experiment created by the University of California, Los Angeles’s suspension of in-person instruction during 2020 and 2021 to explore behavioral plasticity in a trait well-known to vary between urban and non-urban populations of songbirds: territorial aggression (28,21,40,22,41,42,29,43). The dark-eyed junco *(Junco hyemalis;* hereafter, junco) is a recent colonizer of cities in southern California. First observed breeding in San Diego in the early 1980s (44), they have since established multiple populations in other urban centers in the region (45). These urban juncos are behaviorally distinct from wildland counterparts. Protracted breeding seasons and adaptive nesting behavior are prevalent in both the San Diego and Los Angeles populations (25,46,24). Juncos in the San Diego population have a weaker aggression response compared to counterparts in mountain habitats (28,41).

Here we ask: (1) Do junco populations show different amounts of aggression toward conspecific intruders when relieved from frequent encounters with humans? (2) Do individual birds show a plastic aggression response when comparing measurements from before and during the lockdown? With humans temporarily gone from their territories, we predicted that urban juncos would become more aggressive, as the level of human disturbance would more closely resemble that of a wildland habitat. However, our expectation of increased aggression during the anthropause was entirely wrong.

## Results

### Shifts in expression of aggressive behaviors

Our results show that territorial male juncos were less physically aggressive during 2021 compared to 2019 (Fig. 1, Table 1). A principal component analysis (PCA) run on eight measured aggressive behaviors (Table 2) produced three components with an eigenvalue greater than 1, which together explained 78.3% of variance in the data. PC1 accounted for 44.4% of the variance and represented physically aggressive behaviors (Table 3). Its strongest positive associations were with duration of time the focal bird spent at 1–3m distance (0.45) and duration at 0–1m distance (0.43) relative to the model “intruder”, and its strongest negative associations were with nearest distance (−0.46) and latency to 3m distance (−0.43). A low PC1 score describes a bird that did not quickly or closely approach the center of the stage and spent less time within the bounds of the stage: i.e., a bird that was hesitant or unwilling to physically engage with an intruder. PC2 accounted for 18.7% of the variance and represented vocally aggressive behaviors. It had a strong positive correlation with song duration (0.56) and a strong negative correlation with song latency (−0.53). A bird with a low PC2 score waited longer to begin singing and spent less total time singing during the trial. PC3 accounted for 15.2% of the variance. This component had strong associations with number of flyovers (0.76) and number of chip calls (0.58). These behaviors are potentially less risky or energy-intensive compared to the behaviors associated with other aggression components. A t-test comparing PC1 scores from 2019 and 2021 showed that the mean scores across years were not equal (p < 0.001). Mean PC3 scores across years were also not equal (p < 0.001). A t-test comparing PC2 scores across years found no significant difference in means (p = 0.443) (Fig. 1). Further analyses were conducted with physical aggression scores (PC1) only, as these scores explained the most variance in the data and showed the most dramatic behavioral shift.

**Fig. 1:**
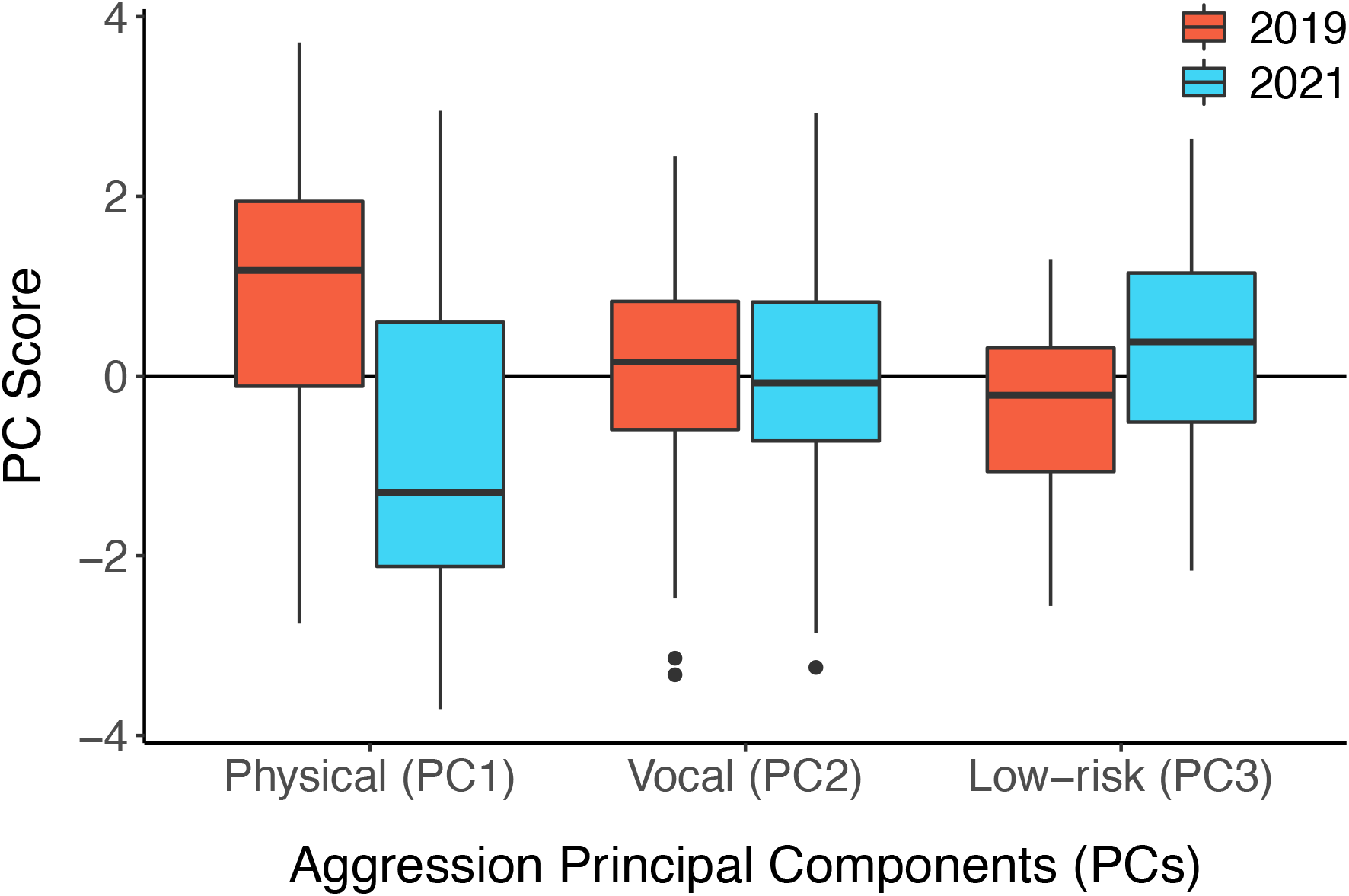
Birds showed significant shifts in two of the top three aggression components when comparing behavior between years 2019 and 2021. PC1 represents physical aggression; PC2 represents vocal aggression; PC3 represents lower-risk behaviors that birds may perform in place of the behaviors associated with physical aggression (PC1). Physical aggression was significantly lower and low-risk aggression was significantly higher during 2021, during the anthropause. Bold lines represent median values, and boxes represent interquartile ranges.

**Table 1:**
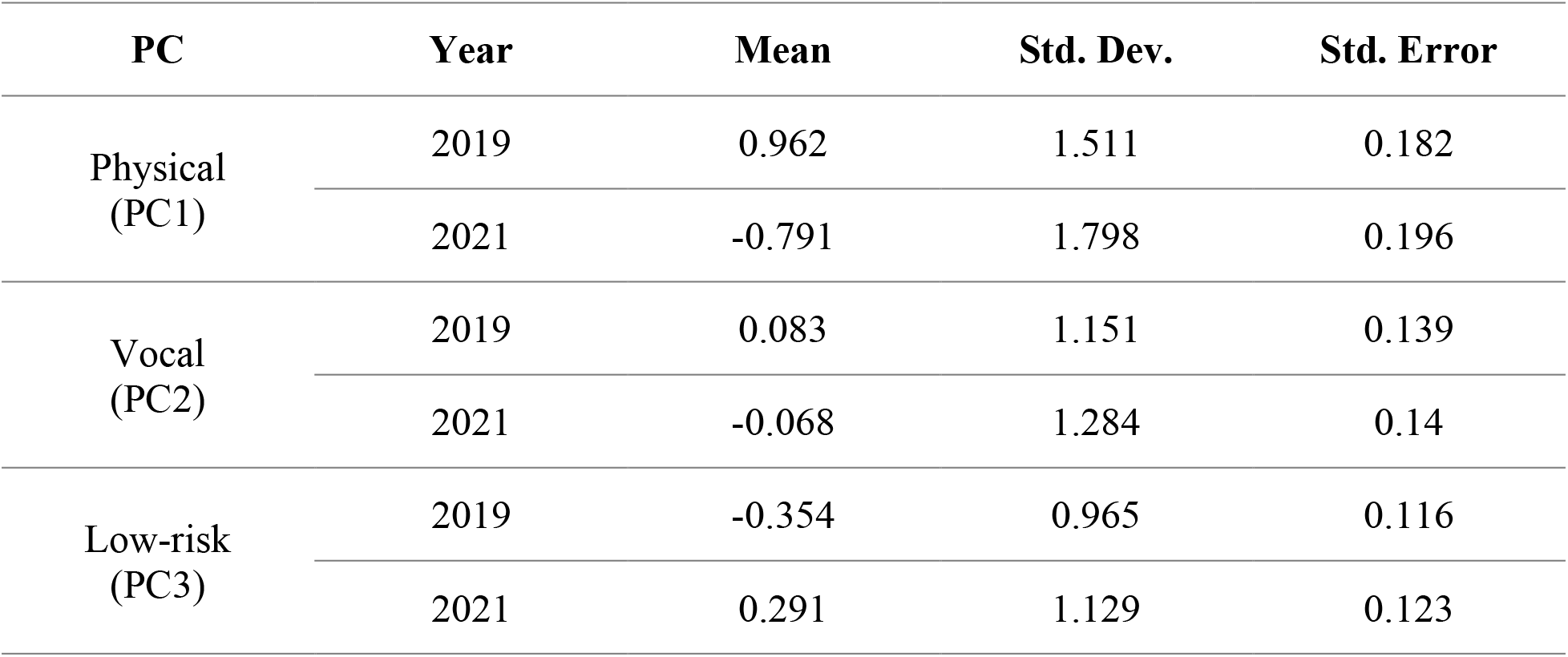
Summary statistics for the top three principal components of aggression. Mean physical aggression scores (PC1) and mean low-risk aggression scores (PC3) were significantly different between 2019 and 2021. There was no significant shift in vocal aggression (PC2).

| PC | Year | Mean | Std. Dev. | Std. Error |
| --- | --- | --- | --- | --- |
| Physical<br>(PC1) | 2019 | 0.962 | 1.511 | 0.182 |
|  | 2021 | -0.791 | 1.798 | 0.196 |
| Vocal<br>(PC2) | 2019 | 0.083 | 1.151 | 0.139 |
|  | 2021 | -0.068 | 1.284 | 0.14 |
| Low-risk<br>(PC3) | 2019 | -0.354 | 0.965 | 0.116 |
|  | 2021 | 0.291 | 1.129 | 0.123 |

**Table 2:** Aggressive behaviors measured during simulated territorial intrusions and summary statistics. We analyzed eight measurements of aggressive behavior recorded during simulated territorial intrusions performed in 2019 and 2021.

| Behaviors | Units | Range | Mean<br>(2019) | Std. Dev.<br>(2019) | Mean<br>(2021) | Std. Dev.<br>(2021) |
| --- | --- | --- | --- | --- | --- | --- |
| Nearest Distance | meters | 0–10 | 1.11 | 1.1 | 2.66 | 2.08 |
| Flyovers | count | 0–39 | 5.55 | 5.59 | 8.82 | 10.59 |
| Chips | count | 0–527 | 15.03 | 27.77 | 67.67 | 95.61 |
| Latency to 3m | seconds | 14–900 | 255.99 | 239.78 | 548.07 | 353.24 |
| Duration 1-3m | seconds | 0–798 | 186.03 | 191.7 | 51.45 | 105.11 |
| Duration 0-1m | seconds | 0–798 | 78.7 | 159.44 | 30.02 | 92.86 |
| Song Latency | seconds | 18–900 | 298.23 | 247.34 | 424.81 | 296.19 |
| Song Duration | seconds | 0–870 | 483.36 | 274.58 | 278.65 | 248.64 |

**Table 3:** Behaviors associated with the top three principal components of aggression. The strongest loadings for each principal component are highlighted. PC1 represents behaviors related to physical approach on the ground (red); PC2 represents singing behaviors (purple); PC3 represents behaviors that may be less risky than those associated with PC1 (blue).

| Behaviors | Physical (PC1) | Vocal (PC2) | Low-risk (PC3) |
| --- | --- | --- | --- |
| Nearest Distance | **-0.463** | 0.277 | 0.019 |
| Latency to 3m | **-0.433** | 0.05 | -0.042 |
| Duration 1–3m | **0.45** | -0.257 | -0.116 |
| Duration 0–1m | **0.427** | -0.248 | -0.111 |
| Song Latency | -0.272 | **-0.594** | -0.1 |
| Song Duration | 0.263 | **0.565** | 0.219 |
| Flyovers | 0.166 | -0.057 | **0.757** |
| Chips | -0.206 | -0.344 | **0.584** |

### Aggression levels of individual birds across both years

Thirteen color-banded, resident birds were measured at UCLA in both 2019 and 2021. A t-test comparing the physical aggression scores from 2019 and 2021 for these individuals showed that the mean level of physical aggression was significantly different across years (p < 0.001) (Fig. 2a). The physical aggression score for 11 out of 13 birds decreased in 2021, during the anthropause (Fig. 2b). In two birds, the level of physical aggression increased; however, the magnitude of the mean change in the 11 birds whose scores decreased (mean = -2.97) was nearly three-fold that of birds whose scores increased (mean = 1.03).

**Fig. 2:**
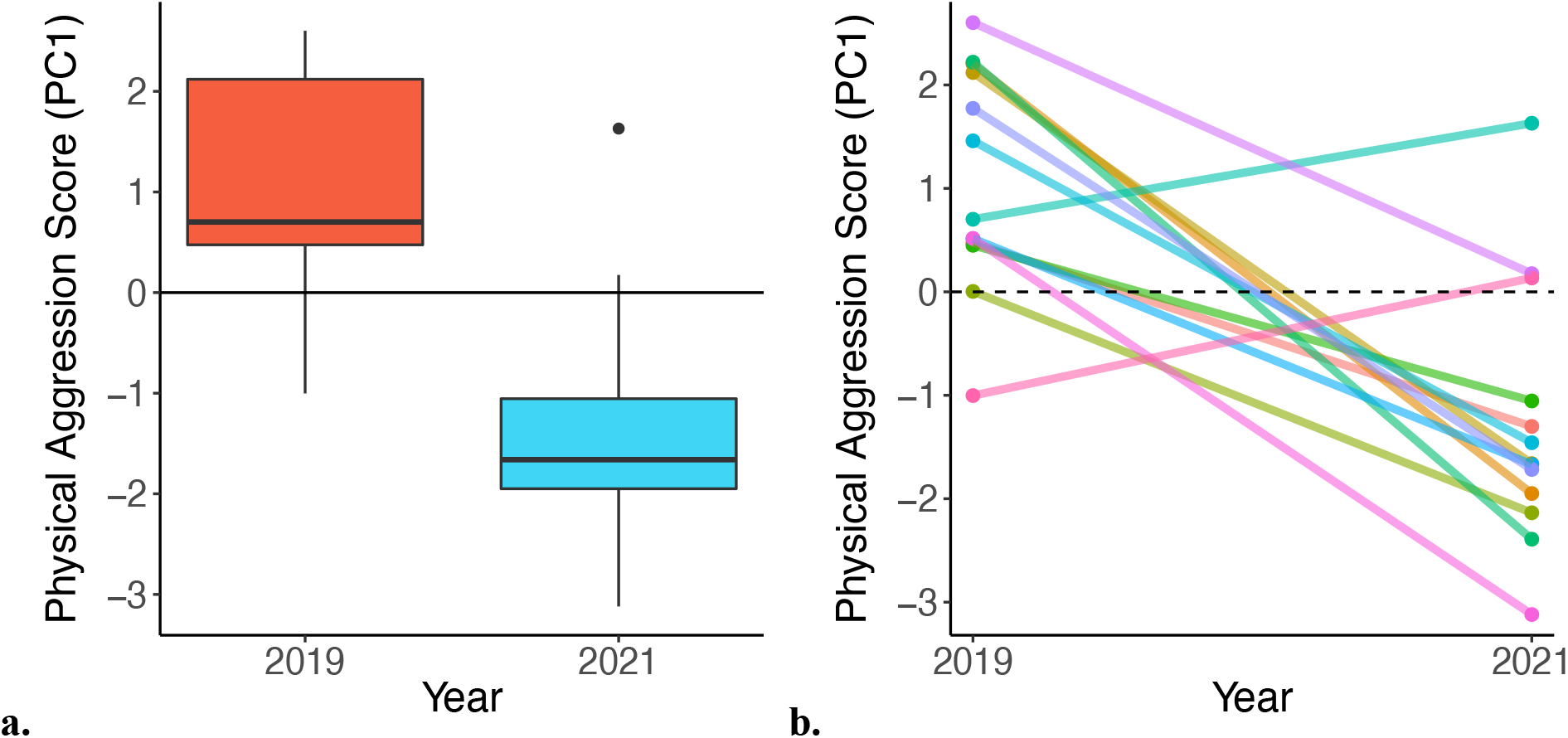
Physical aggression scores (PC1) decreased from 2019 to 2021 in individual birds that were measured in both years. (a) Thirteen individual birds were measured in both 2019 and 2021. Among these birds, the mean physical aggression score (PC1) significantly decreased in 2021. Bold lines represent median values, and boxes represent interquartile ranges. 2021 had one outlier in the data, represented by the black dot. (b) Physical aggression decreased in 11 out of 13 birds. For birds measured multiple times in one year, scores in (b) represent an average of all measurements taken in that year.

### Effect of age on aggression

Our dataset contained 47 observations in which the age of the bird was known. To control for the effect of age on physical aggression level, we used a generalized linear model on the subset of birds with known ages (Table 4). The main effect of age on physical aggression score (PC1) was not significant (p = 0.30), while the main effect of year was (p < 0.001). Thus, we concluded that the difference in human disturbance between years had a significant influence on aggression level, even when controlling for age.

**Table 4:**
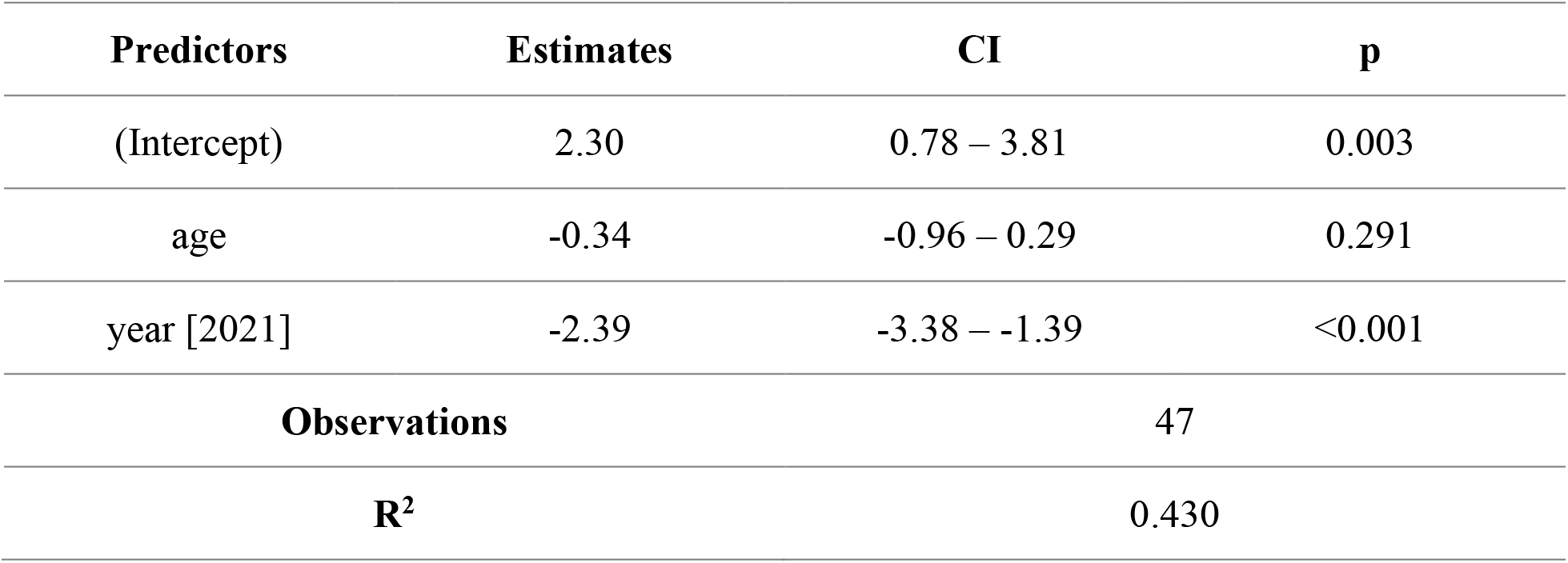
The year explained shifts in physical aggression when controlling for the effect of age. We used a generalized linear model (GLM) to control for the effect of age on physical aggression among the subset of 47 birds whose ages were known. We found that the main effect of year was significant and accounted for the decrease in physical aggression score (PC1), while the main effect of age on physical aggression was not significant.

| Predictors | Estimates | CI | p |
| --- | --- | --- | --- |
| (Intercept) | 2.30 | 0.78 – 3.81 | 0.003 |
| age | -0.34 | -0.96 – 0.29 | 0.291 |
| year [2021] | -2.39 | -3.38 – -1.39 | <0.001 |
| Observations |  | 47 |  |
| R<sup>2</sup> |  | 0.430 |  |

### Pedestrian point counts

A t-test showed that the number of people on campus was significantly less in 2021 compared to 2022, a year with typical pedestrian traffic (p < 0.001). We observed a nearly seven-fold increase in average pedestrian traffic across campus in 2022 (Fig. 3, Table 5). In 2022, we conducted 156 surveys and did not have a single survey in which no humans were present; in 2021, 72 out of 312 surveys (23%) had no humans present. Disturbance from pet dogs and large motor vehicles occurred rarely in both years (< 1 per survey) and did not significantly change between years.

**Fig. 3:**
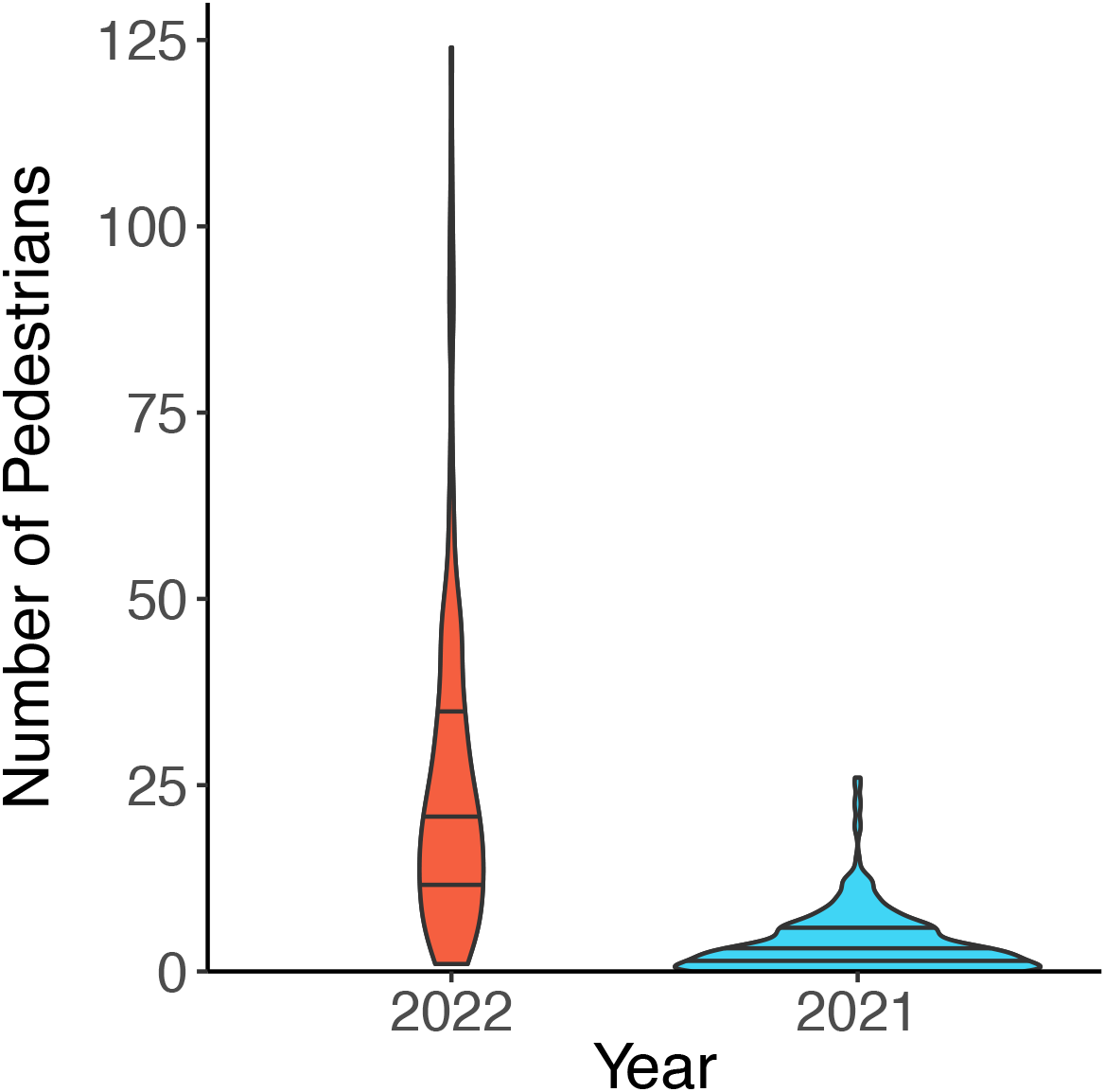
Pedestrian traffic was significantly lower in pandemic lockdown year 2021 compared to a non-lockdown year 2022. Violin plots show numbers of pedestrians counted during each year’s surveys. Wider areas represent more surveys where pedestrian counts were similar to each other. Lines represent quartiles. About one quarter of surveys conducted in 2021 had no people present. All surveys in 2022 had at least one person present. There was a near seven-fold increase in mean pedestrian traffic in the year 2022 compared to 2021. Because all campus restrictions were lifted and all classes were taught in-person starting in March 2022, we expect pedestrian traffic in spring of 2022 is approximately similar to that of 2019.

**Table 5:** Summary statistics for pedestrian surveys. Mean pedestrian traffic on campus was significantly lower during 2021 compared to a typical year. To quantify human disturbance, we counted the number of pedestrians passing through 13 outdoor locations approximately evenly spaced around the UCLA campus. All surveys were two minutes in length. As all restrictions were lifted and all classes were held in-person starting in March 2022, conditions in 2022 are likely similar to those in 2019.

| Year | Mean | Median | Std. Dev. | Minimum | Maximum | Total Surveys |
| --- | --- | --- | --- | --- | --- | --- |
| 2022 | 25.3 | 20 | 22.1 | 1 | 124 | 156 |
| 2021 | 3.6 | 3 | 4.1 | 0 | 26 | 312 |

## Discussion

Resident juncos in urban Los Angeles defended territory less aggressively during the anthropause in 2021 compared to 2019, a typical non-lockdown year. Reducing territorial aggression may benefit individuals because defending territory less aggressively allows for allocation of energy to other tasks, like parental provisioning (47). Less aggression may also reduce the risk of confrontations with rivals that lead to injury (48). Our analysis identified three categories of territorial aggressive behavior: physical aggression (PC1), vocal aggression (PC2), and a third category that may represent lower-risk territorial behaviors (PC3). We saw the most significant change in physical aggression: birds in 2021 were much less likely to engage in risky, physically aggressive behavior, such as approaching the intruder on the ground. We saw no difference between years in vocal aggression; singing behaviors likely carry little risk of injury because birds can perform them from off-ground perches. The small but significant change in PC3 in 2021 may indicate that birds replaced the riskiest form of physical aggression—approach on the ground—with a less confrontational form—flyover—to reduce energy expenditure or the risk of injury.

Crucially, a subset of thirteen individual birds shifted their responses and showed significantly decreased aggression in 2021 compared to 2019, indicating that territorial aggression is a plastic behavior that adult birds can modify to react to changes in their environment. Our results from individuals allow us to rule out other potential mechanisms for this population-wide behavioral shift, such as reduced human disturbance allowing birds expressing a fixed, lower level of aggression to become more prevalent in the population (49,50).

Our prediction that juncos would act more aggressively in the absence of human disturbance was the opposite of what we observed. We originally expected that lower levels of noise and distraction generated by human activity would improve an individual’s ability to perceive its neighbors, leading to more aggressive territorial conflicts (51,52). Furthermore, other experiments conducted during the anthropause showed urban birds behaving more similarly to wildland populations (36,38), and wildland juncos are more aggressive toward intruders than their urban counterparts (28,41).

Yet, when we consider aggression levels in light of resource availability, our results become more understandable. Territorial aggressive behavior often differs between urban and wildland songbird populations (43), and can serve as a means to understand how urban individuals experience resource availability in a novel environment (53–55). If resources are scarce or patchily distributed, individuals may increase aggression to defend valuable territories from intruders; if resources are evenly spread, individuals may expend less energy on territory defense and more on parental provisioning or other tasks (47). Because urban environments are characterized in part by high habitat fragmentation and a mixture of native and exotic vegetation (1), this spatial and biotic heterogeneity likely equates to uneven resource availability for many songbird species. Urban bird populations that rely on specific vegetation for food or nesting sites may have their distributions or population density limited by the presence of suitable habitat within the urban matrix. For example, increased territorial aggression in urban curve-billed thrashers *(Toxostoma curvirostre)* and Abert’s towhees *(Melozone aberti)* was strongly correlated with the presence of desert vegetation preferred for nesting (40). Similarly, increased territorial aggression in an urban population of song sparrows *(Melospiza melodia)* may be driven by lower availability of breeding sites (29).

However, if resources are overly abundant across the landscape, birds may lose motivation to defend territories, because expending energy to exclude rivals offers no additional resource availability (55). Urban juncos could be flush with resources compared to similar species, thus explaining their shift towards less territorial aggression while other songbirds, like the song sparrow, become consistently more aggressive in urban areas (43). In contrast to species that need specialized nesting habitat, the adaptive nesting behavior of urban juncos unlocks abundant potential nesting sites above ground and on artificial structures (46,24), possibly reducing intraspecific competition for these sites. During the pandemic lockdown, foraging and nesting sites previously disturbed by humans likely became available for juncos to use continuously, causing a sudden increase of resources in the environment and thus a decrease in territorial behavior.

Another and complementary explanation for decreased aggression in a newly decreased human-disturbance environment is that juncos in 2021 became less vigilant when relieved from frequent human disturbance. Birds may consider humans to be predators because of how often humans unwittingly approach them and occupy valuable foraging space (56,57). Having to actively avoid humans may strongly encourage vigilance behavior in adult birds. This in turn draws investment away from foraging and provisioning chicks and promotes avoidance of areas of the landscape with high predator activity (57–59). Juncos perceiving less predation risk from humans may be less vigilant toward competitors as well if they shift energetic investment to parental care.

Because vigilance and nest defense behaviors are associated with trade-offs in parental care (58,59), limiting human disturbance of urban bird populations long-term could potentially improve the overall health and survival of chicks. This trade-off was evident in a pandemic lockdown-era study which measured the body condition of nestling urban great tits *(Parus major)* in urban parks of two cities during the lockdowns: one in which community park attendance increased, and one in which it decreased. Nestling great tits in busier parks had worse body condition, indicating that adults invested less energy in parental provisioning compared to the prior year when human disturbance was not as intense. However, the body condition of chicks in less-attended parks did not improve versus a typical year, suggesting that short-term relief from human disturbance is not enough to confer lasting benefits to urban birds (8).

Adaptive nesting behavior has been seen in both in the Los Angeles population studied here and another, independently established population in La Jolla, California, on the University of California, San Diego campus. For example, urban juncos are more likely to build off-ground nests and use artificial structures as nesting sites in both the San Diego and Los Angeles populations (46,24). In San Diego, 20% of females built off-ground nests, compared to wildland populations where the proportion was much lower (60). This behavior is highly adaptive, as eggs laid in off-ground nests were ∼80% more likely to result in a chick hatching and surviving to the following year versus eggs laid in on-ground nests. Individual females built both on-and off-ground nests during the same breeding season, demonstrating plasticity in nest site selection (46). Juncos in Los Angeles also switched between on-ground and off-ground nesting sites within a single breeding season depending on the outcomes of their prior nesting attempts— juncos were more likely to switch strategies when their prior nest failed (24). Additionally, the overall population off-ground nest rate was 38% and correlated with increased fitness.

Phenotypic plasticity in breeding behavior has likely contributed to the overall success of urban junco populations.

We considered several alternative explanations for shifting aggressive behavior in urban juncos, instead of or in addition to the effect of reduced human disturbance: (1) differences in climate, (2) changes in predation levels, and (3) differences in body condition (see Supplementary Information). None of these proved significantly different between a lockdown and non-lockdown year. Because the UCLA campus vegetation is regularly watered and temperature across years was for the most part similar, we do not expect that resource availability from vegetation was meaningfully different. Reduced human activity may have affected the behavior of junco nest predators, but we found no evidence that predation or the overall nest failure rate changed across years. While body condition in either nestling or adult birds did not improve during the anthropause, our results are consistent with another study that showed no changes in the body condition of urban birds when human activity was temporarily reduced (8). We found that reduced human disturbance was the most substantial difference in environmental conditions on the UCLA campus between 2019 and 2021 and thus the most likely driver of our results.

Our finding of a population-wide shift in behavior is consistent with the hypothesis that large-scale lockdowns of humans would affect the behaviors of other urban animals. For example, urban white-crowned sparrows *(Zonotrichia leucophrys)* in the Bay Area of California sang lower amplitude, higher quality songs that better transmitted salient information while the environment was quieter (38). In Catalonia, Spain, several species of urban-dwelling songbirds observed by community scientists were more detectable in early morning during lockdowns, exhibiting temporal singing patterns that shifted to resemble those of birds in wildland environments (36). Both studies show that urban bird populations maintain variation in singing behavior suitable for rapidly changing environments like cities.

The COVID-19 pandemic and associated lockdowns created an unprecedented opportunity to study the behavior of urban wildlife in the absence or marked decrease of human disturbance, a stressor that strongly shapes the behavior and distribution of wildlife (61,5,62). By taking advantage of this natural experiment, we identified significant shifts in territorial aggression in urban juncos. Our results support the idea that maintenance of behavioral plasticity is critical to the success of newly established wildlife populations (25,30) and underscores the crucial role phenotypic plasticity plays in the success of birds in urban areas.

## Materials and Methods

### Dates and location

Birds measured were residents of the University of California, Los Angeles (hereafter, UCLA) campus in Westwood, Los Angeles, California. UCLA’s campus has a mild, costal climate with abundant trees, shrubs, and foliage, both ornamental and native, as well as a high density of tall buildings and paved ground surfaces interspersed with grassy lawns. In the spring of 2019, UCLA had 41,569 enrolled students and 36,588 faculty and staff visiting campus regularly. In March 2020, UCLA cancelled all in-person instruction and implemented protocols that drastically reduced the number of faculty, staff, students, and visitors on campus. These restrictions to campus traffic continued throughout 2021, with some restrictions partially lifted in September 2021 and all restrictions lifted in March 2022.

### Quantifying human disturbance

Twice weekly from 11 May to 28 July in 2021 and from 1 April to 17 May in 2022, pedestrian point count surveys took place at 13 locations distributed approximately evenly across the UCLA campus. Each week, one set of surveys was completed in the morning between 8:30–11am, and the other set in the afternoon between 12–3pm. For each point count, a single observer tallied all people passing through their line of sight for two minutes. When applicable, the observer also tallied pet dogs and large motor vehicles, but these disturbance types did not occur at all locations. We completed 312 surveys in 2021 and 156 surveys in 2022. Because all campus restrictions were lifted and all classes were held in-person starting in March 2022, we expect that 2022 survey results represent a roughly similar amount of human disturbance as 2019.

### Trapping and tracking individuals

Adult birds were trapped with mist nets using a recording of territorial songs as an audio lure. Each captured bird was affixed with one metal USGS band and a unique combination of three colored bands for identification from a distance. Sex and age (whenever possible) were determined at capture by primary sex characteristics (brood patch or cloacal protrusion), plumage, and molt limit. Length of the tarsus (to the nearest 0.1mm) and body weight (to the nearest 0.1g) were taken to calculate overall body condition. Dimensions of the bill, wing length, and tail length were also recorded, and a blood sample was taken. Territories of adult birds were determined through marking locations of singing perches on a map, followed by observations of nest building behavior and mapping of nest locations as the breeding season progressed. Found nests were monitored until their final outcomes were determined. Nestling juncos were banded, measured, and bled at the age of seven days.

### Simulated territorial intrusions

Aggression in songbirds is often measured by using simulated territorial intrusions, a method that uses a decoy model or captive live specimen, pre-recorded audio of a competitor’s song, or both, to stimulate a territorial response in the target wild bird (63–65). An observer then records and analyzes specific behaviors to quantify the level of aggression of each individual. This method generates a standardized measurement of a multifaceted behavior with many modes of expression across individuals within a population.

Birds were measured during their breeding season between March to July 2019 and March to July 2021. Urban resident juncos have a breeding season that is about twice as long as that of typical, migratory juncos (25,24). The dataset we analyzed comprises 49 unique male birds in the 2019 breeding season and 76 unique males in 2021; including repeated trials, our totals are 69 and 84 measurements per year respectively. In addition, five trials were conducted in the spring of 2020 before the campus lockdown went into effect and research was discontinued in the interest of personal safety and changing campus pandemic regulations. Because of the small sample size, we did not include data from 2020 in our analysis.

### Playback recordings

Three unique, 15-minute audio tracks were compiled using field recordings of junco songs taken on the UCLA campus during the 2018 breeding season. The songs of three randomly selected birds appear on each track. The recordings were made with a Marantz PMD661 solid-state digital recorder and either a Sennheiser ME66 omnidirectional microphone or an Audio-Technica AT815b microphone. The recordings were saved as WAV files with a 44 kHz sampling rate. Each song recording was normalized in Audacity 2.3.0 (66).

### Simulated territorial intrusion procedure

The trial stage was a circle three meters in radius, marked with two 6-meter lengths of nylon cord laid perpendicular, crossing at their centers, and staked to the ground at their ends. Flagging was tied to each length of cord to mark the 1-meter radius inner circle. At the center of the circle was a wireless speaker (NYNE Boost IP67), and atop the speaker was placed a 3d-printed and painted model of a male junco (Fig. 4). The stage was set up in a wide-open area as close as possible to the location the target male was previously observed singing, or as safely possible to an active nest (∼5m away from on-ground nests). Whenever possible, observers chose a location without low vegetation or other perches within the stage area to allow the target bird to move freely and to avoid bias against those trials taking place on paved surfaces.

**Fig. 4:**
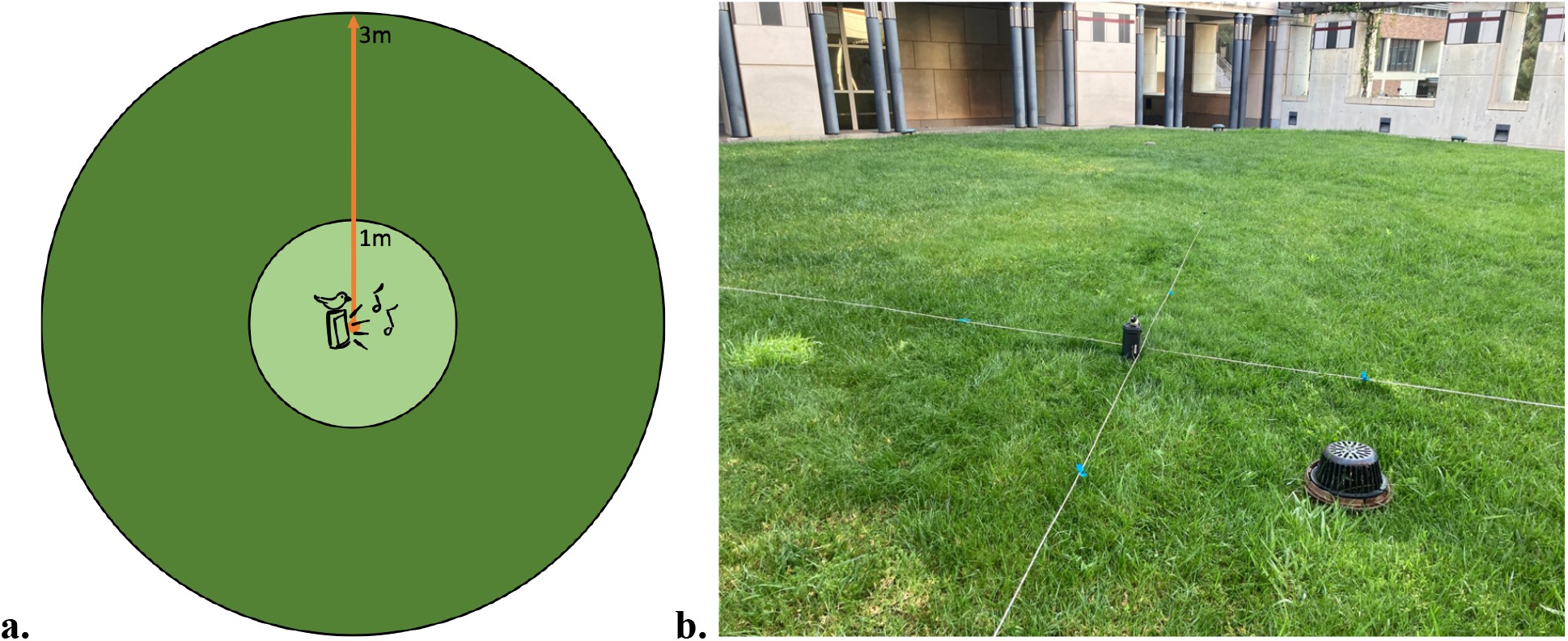
Set-up and dimensions of the simulated territorial intrusion stage. A diagram (a) and a photograph (b) of the simulated territorial intrusion stage. The stage has a radius of 3m, with an inner radius of 1m marked with flagging tape. At the center is a portable speaker and decoy model of a junco.

Each trial was 15 minutes in length. Observers retreated at least 10m from the outer edge of the stage before beginning playback. One randomly selected playback recording was played at maximum volume (L_max_ ∼85dB at 1m) with the same speaker for each trial, though the perceived volume may have fluctuated as structures and ground surfaces are highly variable throughout the UCLA campus. Observers ensured that the selected playback did not contain the song of a neighboring bird to counteract the “dear enemy” effect, as a bird may respond less aggressively to the songs of birds that defend territories neighboring its own (67,68).

During playback, observers recorded the following behaviors: nearest distance to the speaker, number of flyovers (defined as any flight passing directly over the stage), number of chips (a vocalization conveying alarm or agitation), latency to enter the 3m outer radius of the stage, duration spent within 1–3 meters of the speaker, duration spent within 0–1 meters of the speaker, latency to perform a long-range song, and duration of singing (Table 2).

The majority of trials took place between 6am and 11am, with a few exceptions where birds were measured as late as 2pm. Males were not measured within seven days of capture to limit exposure to the audio stimulus. Males tested multiple times throughout the season were given at least two weeks of cool-down between trials. Measurements of the same male taken in the two weeks after a prior measurement were discarded from the analysis to avoid any influence of habituation to the stimulus.

Due to the high density of territories on the UCLA campus, a few trials attracted more than one territorial bird. If multiple birds were present and the observers were confident that they would not misattribute behaviors to the wrong birds, they recorded the behavior of all birds simultaneously. Two competing males were rarely recorded at the same trial (N = 6 in 2019, N = 3 in 2021). Aggressive female birds were recorded in a few instances (N = 4 in 2019, N = 8 in 2021). Data from females were discarded from the final analysis because they express aggression differently than males, e.g. they are much less likely to sing.

### Analysis

All analyses were done with R 4.1.3 (69) and R Studio 2022.2.1 (70).

We analyzed the behavioral measurements collected during simulated territorial intrusions using principal component analysis (PCA). To normalize the distributions before analysis, all variables were log-transformed by adding a constant 1 then taking the natural log. We used two-tailed t-tests to compare each principal component across years. Further analysis of individual birds measured in both years were conducted with physical aggression scores (PC1) only, using a paired t-test. To control for the effect of age on physical aggression level, we used a generalized linear model on the subset of 47 birds with known ages, with physical aggression score (PC1) as the response variable and age and year as predictor variables.

We used a two-tailed t-test to compare the amount of pedestrian traffic on the UCLA campus in 2021 and 2022.

## Acknowledgements

We thank Dr. Morgan Tingley for consultation on the statistical analyses used in this work. We thank Donovan Roediger for comments on the manuscript. We thank the Santa Monica Bay Audubon Society and the San Fernando Valley Audubon Society for funding.

## Supplementary Information

In addition to the analyses presented in the main text, we also considered alternative explanations for differences in aggressive behavior during the pandemic lockdown. We analyzed (1) body condition of nestlings and adults, (2) rates of nest failure and predation, and (3) climate differences across the years 2019 and 2022.

## Results

### Body condition of nestlings and adults

Nestling and adult juncos had similar body condition in both years. T-tests showed no significant difference in mean body condition index in either nestlings (p > 0.9) (Supplementary Fig. 1a) or adults (p > 0.9) (Supplementary Fig. 1b) across years, indicating that body condition did not significantly change in the population between 2019 and 2021.

**Supplementary Fig. 1:**
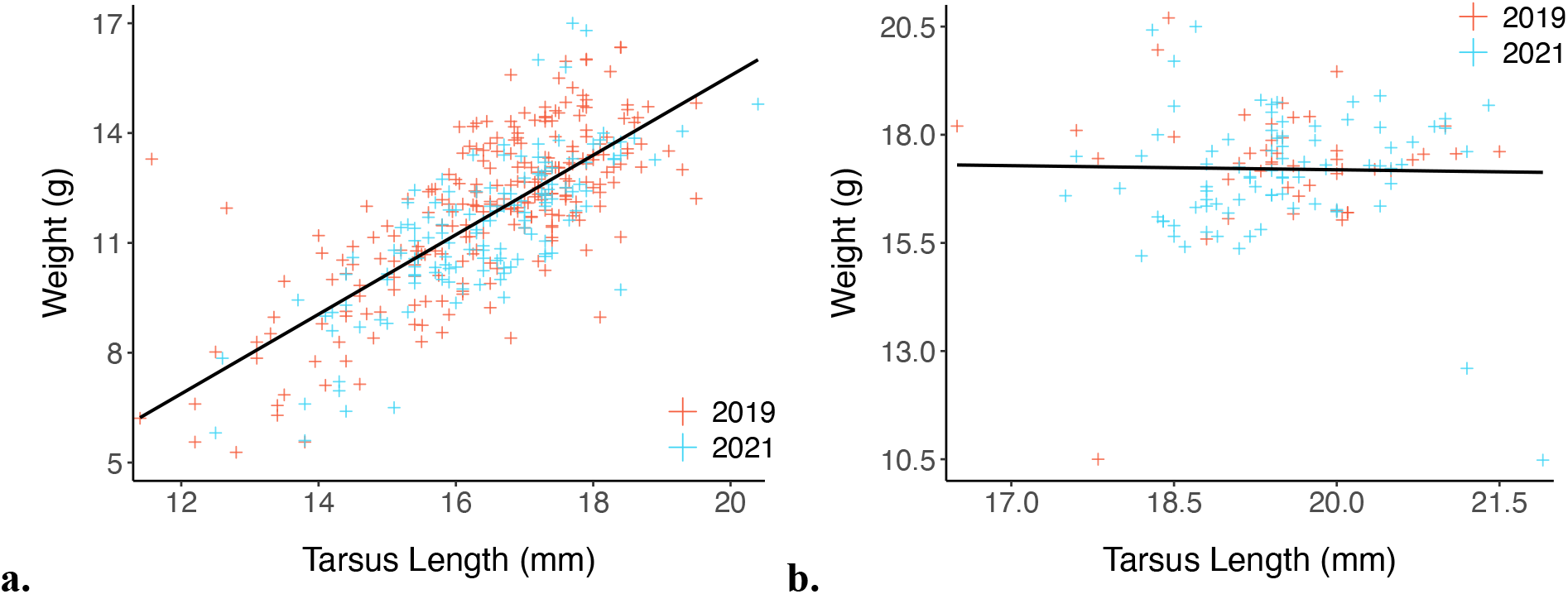
Body condition was not different in either nestlings or adults across years. We used linear regression to calculate body condition scores from tarsus length and body weight in (a) nestling and (b) adult juncos. We found no significant differences when comparing body condition scores (residuals) in either nestlings or adults. Plots show the distribution of individuals (crosses) around the line of best fit (in black).

### Nest Failure and Predation

The nest failure rate was relatively low in the population in both years. In 2019, the rate of nest failure for any reason was 25.1% (45 of 179 nests); in 2021, the rate was 26.1% (41 of 157 nests). Of these nest failures, 8 (4.5%) in 2019 and 4 (2.5%) in 2021 were directly attributed to predation. Using a logistic regression model, we found no significant difference in the overall failure rate across years (p = 0.026). Neither did we find a significant difference in predation rate (p = 0.291).

### Climate

Average daily temperatures at UCLA were mostly similar between 2019 and 2021. T-tests showed only May had a significant difference in mean average daily temperature between years (Bonferroni-corrected p < 0.001) (Supplementary Fig. 2a). Total precipitation at UCLA was very low in 2021, with four out of five months having less than half a centimeter of rainfall (April– July), while 2019 had two months (March and May) with several centimeters of total precipitation (Supplementary Fig. 2b). However, the UCLA campus is extensively irrigated, and lawns were watered regularly in both years.

## Methods

**Supplementary Fig. 2:**
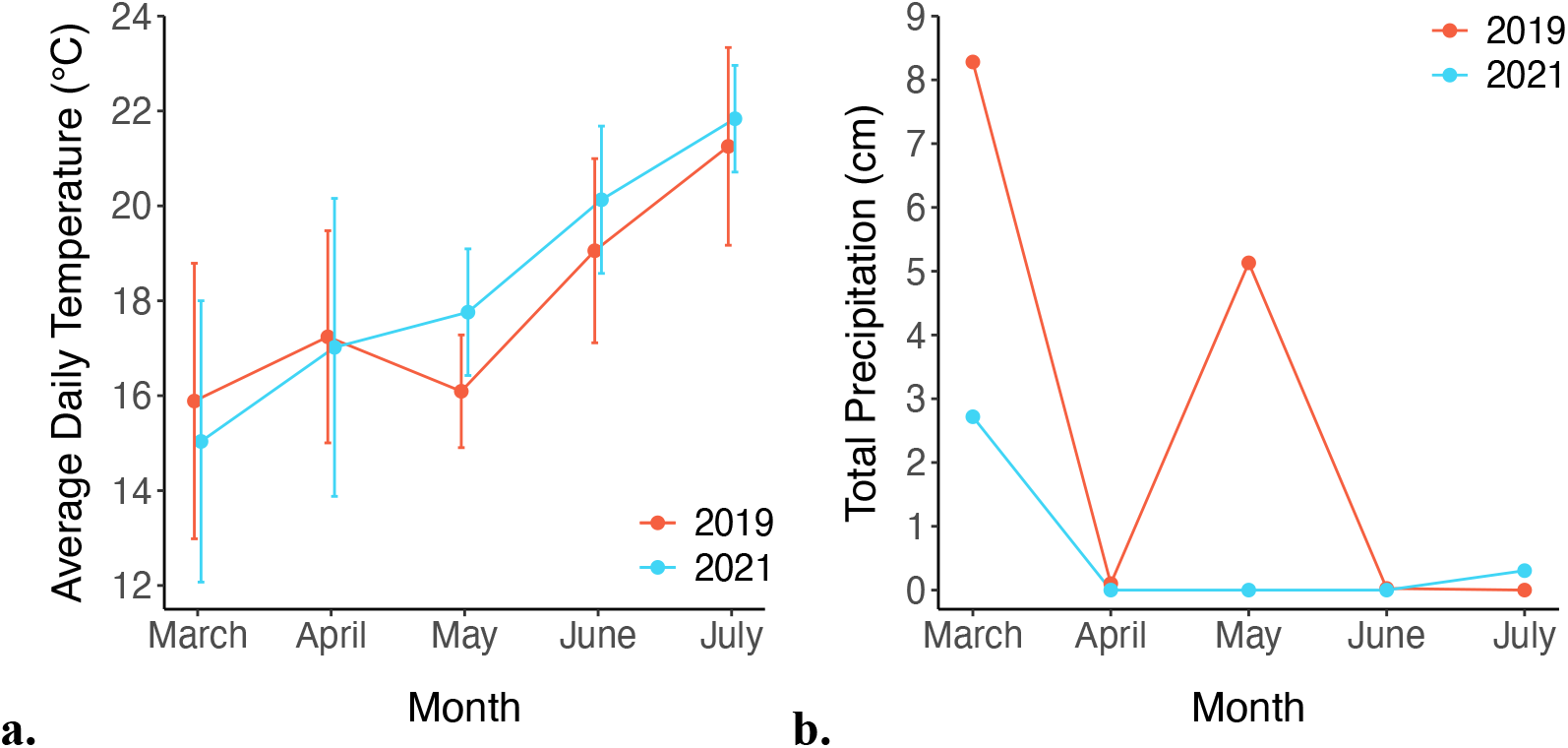
Average daily temperature was mostly similar between years while total precipitation varied. (a) Only the month of May had a significant difference in mean average daily temperature between years. Error bars represent standard deviations. (b) While precipitation was more variable between years, with 2021 being exceptionally dry, we expect the effect on vegetation was minimal because of extensive irrigation and regular watering on the UCLA campus.

To calculate a body condition index, we used the residuals from a linear regression analysis of body weight (a measure related resource availability) on tarsus length (a heritable indicator of body size) in nestling and adult birds. To remove outliers from the data before regression, we calculated z-scores for tarsus and weight measurements for chicks and adults and excluded those outside of three standard deviations from the mean for either measurement. Our analysis included 381 chicks (N = 243 in 2019, N = 138 in 2021) and 127 adults (N = 45 in 2019, N = 82 in 2021).

To determine if nest failure rates changed between years, we used a logistic regression model with nest fate as the response variable and year as the predictor variable using the recorded outcomes of all found nests in 2019 and 2021.

We used a series of two-tailed t-tests to compare mean average daily temperature for the five months trials were performed (March–July) in 2019 and 2021. We obtained past average daily temperature and monthly total precipitation data from the National Weather Service UCLA station.

